# Dorsal Striatum Parvalbumin interneurons translatome unveiled

**DOI:** 10.1101/2025.06.16.659856

**Authors:** Claire Naon, Laia Castell, Steeve Thirard, Maria Moreno, Stéphanie Rialle, Eva Goetz, Eloi Casals, Angelina Rogliardo, Marta Gut, Anna Esteve-Codina, Albert Quintana, Federica Bertaso, Emmanuel Valjent, Laura Cutando

**Affiliations:** INM, University Montpellier, Inserm, 34091 Montpellier, France; Department of Neuroscience, Northwestern University Feinberg School of Medicine, Chicago, IL 60611, USA; IGF, University Montpellier, CNRS, Inserm, 34094 Montpellier, France; MGX-Montpellier GenomiX, University Montpellier, CNRS, Inserm, Montpellier, France; Centro Nacional de Análisis Genómico (CNAG), Baldiri Reixac 4, 08028 Barcelona, Spain; Universitat de Barcelona (UB), Barcelona, Spain; Institut de Neurociències, Universitat Autònoma de Barcelona, Bellaterra, Spain; Departament de Biologia Cellular, Fisiologia i Immunologia, Universitat Autònoma de Barcelona, Barcelona, Spain

## Abstract

Parvalbumin (PV) interneurons in the dorsal striatum (DS) are fast-spiking GABAergic cells critical for feedforward inhibition and synaptic integration within basal ganglia circuits. Despite their well-characterized electrophysiological roles, their molecular identity remains incompletely defined. Using the Ribotag approach in *Pvalb-Cre* mice, we profiled the translatome of DS PV interneurons and identified over 2,700 transcripts significantly enriched (fold-change > 1.5) in this population. Our data validate established PV markers and reveal a distinct molecular signature of DS PV neurons compared to PV interneurons from the nucleus accumbens. Gene ontology analyses highlight prominent expression of genes related to extracellular matrix components, cell adhesion molecules, synaptic organization, ion channels, and neurotransmitter receptors, particularly those mediating glutamatergic and GABAergic signaling. Notably, perineuronal net markers were robustly expressed in DS PV interneurons and confirmed by immunofluorescence. Transcriptomic analysis of DS PV neurons following repeated d-amphetamine exposure identified *Gm20683* as the only differentially expressed transcript between treated groups. Furthermore, RNAseq analysis of mice subjected to an operant behavior paradigm with two types of food reward (high-palatable diet or standard chow) identified over 1,000 and 100 genes enriched in DS PV neurons from standard and high-palatable masters, respectively. These findings provide a comprehensive molecular profile of DS PV interneurons, distinguishing them from other striatal PV populations, and reveal specific gene expression changes associated with psychostimulant exposure and reward-driven behaviors. Our findings deepen insight into the molecular mechanisms of PV interneuron activity in striatal circuits and their potential roles in neuropsychiatric, motor and reward-related disorders.

## Introduction

The dorsal striatum (DS) is the primary input nucleus of the basal ganglia, alongside the ventral striatum. It receives converging excitatory inputs from both the cortex and the thalamus (Graybiel and Grafton, 2015; Hintiryan et al., 2016; Hunnicutt et al., 2016; Pan et al., 2010; Smith and Bolam, 1990; Tisch et al., 2004). These inputs are subsequently processed and converted into an inhibitory signal projecting toward the basal ganglia output nuclei. Based on the targeted output regions, two main GABAergic projection pathways are typically described: the striatonigral (direct) pathway, which projects to the substantia nigra pars reticula (SNr) and the internal pallidum (GPi), and the striatopallidal (indirect) pathway, which projects to the external pallidum (GPe) (Oh et al., 2014). The direct and indirect striatal projection neurons (dSPNs and iSPNs) define these downstream pathways respectively (Gerfen et al., 2013; Kreitzer and Malenka, 2008; Marsden and Obeso, 1994). SPNs constitute approximately 95% of all striatal neurons (Tisch et al., 2004), the remaining 5% being interneurons that form a highly organized and hierarchical network, essential for efficient input processing and proper striatum function (Graveland and DiFiglia, 1985; Tepper et al., 2018).

The population of dorsal striatal interneurons can be classified into either cholinergic or GABAergic subtypes which comprise four major classes based on their molecular and firing properties. These clusters include interneurons expressing parvalbumin (PV), neuropeptide Y, tyrosine hydroxylase and calretinin (Kreitzer, 2009; Tepper et al., 2018; Tepper and Bolam, 2004). Among them, PV interneurons, also described as fast-spiking interneurons (FSIs), represent the main population of GABAergic interneurons (Tepper et al., 2018). By receiving converging excitatory inputs, they mediate rapid feedforward inhibition of SPNs, thereby modulating the striatal plasticity necessary for the proper integration of excitatory information by SPNs (Arias-García et al., 2018; Choi et al., 2019; Duhne et al., 2021; Gage et al., 2010; Gittis et al., 2010; Mallet et al., 2005; Owen et al., 2018; Planert et al., 2013; Roberts et al., 2019; Sciamanna et al., 2015; Tepper and Bolam, 2004). Importantly, disruption of PV activity has been associated with core motor symptoms of neurological and neurodevelopmental disorders including obsessive-compulsive disorder (OCD), dyskinesias, Tourette syndrome, motor stereotypies in autism spectrum disorder (ASD) and schizophrenia (Burguière et al., 2013; Gittis et al., 2011; Mondragón-González et al., 2024; Zhou et al., 2021). Surprisingly, although DS PV interneurons have been extensively studied at the functional level, insight related to their molecular profile remain limited to a restricted number of genes identified using single-cell transcriptomics (Muñoz-Manchado et al., 2018; Saunders et al., 2018). The present study aimed to establish the translatome of DS PV interneurons using the Ribotag approach, allowing cell type-specific gene expression profiling (Puighermanal et al., 2020; Sanz et al., 2009). We generated *Pvalb-Ribotag* mice and identified over 2.700 transcripts enriched in DS PV interneurons. Additionally, we investigated whether repeated exposure to d-amphetamine, as well as, food-seeking behaviors (standard vs high palatable food) and contingency (master vs yoked mice) impacted gene expression in DS PV interneurons

## Materials and Methods

### Animals

All animal procedures were conducted in accordance with the guidelines of the French Agriculture and Forestry Ministry for handling animals (authorization number/license B34-172-41) and approved by the relevant local and national ethics committees (authorization APAFIS#38912). Animals were housed in groups of 2 to 5 per cage under standardized conditions with a 12-hour light/dark cycle, *ad libitum* food and water, stable temperature (22 ± 2°C) and controlled humidity (55 ± 10%). Mouse strains used for immunofluorescence included *Pvalb-ChR2-Ribotag* mice (n = 2), *Pvalb-ChR2* mice (n = 3), *Pvalb-Ribotag* mice (n = 6) and *Pvalb-Ai14* mice (n = 3). *Pvalb-Ribotag* mice were used for low-input RNA-sequencing (RNA-seq). The experimental groups included: saline (n = 7), D-amphetamine (n = 8), highly palatable food master (n = 6), highly palatable food yoked (n = 6), standard food master (n = 6), and standard food yoked (n = 6). For each RNAseq sample, striatal tissue from two to three mice were pooled.

### Drugs and treatments

(+)-α-Methylphenethylamine [D-amphetamine (D-amph)] sulfate salt (5 mg/kg) from Tocris was dissolved in 0.9% (w/v) NaCl (saline) and injected intraperitoneally (i.p) in a volume of 10 ml/kg. Mice were administered with d-amphetamine (5 mg/kg) during 5 days and euthanized 3 days after the last injection.

### Immunofluorescence

Free-floating sections (30 μm) of the dorsal striatum were prepared as previously described (Cutando et al., 2021). On day 1, slices were washed for 10 min in PBS (3x), incubated 15 min in 0.2% Triton X-100 in PBS before incubation overnight at 4°C with primary antibodies including chicken anti-GFP (1:500, Invitrogen, #A10262), rat anti-HA (1:500, Roche, #11867431001), goat anti-PV (1:500, Swant, #PVG-213), rabbit anti-PV (1:1000, Swant, #PV25), mouse anti-RFP (1:1000, MBL, #M155-3), rabbit anti-TTF1 (1:500, Santa-Cruz, #sc-13040) and N-acetylgalactosamine-binding Wisteria floribunda agglutinin (WFA, 1:1000, Sigma, #L1516). Non-specific binding was blocked with 10% Normal Donkey Serum (NDS) in PBS for 2 hours for WFA staining. On day 2, slices were rinsed in PBS and incubated 45 min with various combination of the following secondary antibodies: goat Alexa Fluor 488-coupled anti-chicken (1:500, Jackson ImmunoResearch, #103-545-155), donkey Alexa Fluor 488-coupled anti-goat (1:500, Jackson ImmunoResearch, #103-545-003), donkey Alexa Fluor 647-coupled anti-goat (1:500, Jackson ImmunoResearch, #103-605-003), donkey Alexa Fluor 594-coupled anti-mouse (1:500, Jackson ImmunoResearch, #103-585-150), goat Alexa Fluor 594-coupled anti-mouse (1:500, Jackson ImmunoResearch, #103-585-003), goat Alexa Fluor 488-coupled anti-rabbit (1:500, Invitrogen, #A11034), goat Cy3-coupled anti-rabbit (1:500, Jackson ImmunoResearch, #111-165-003), donkey Alexa Fluor 647-coupled anti-rat (1:500, Jackson ImmunoResearch, #712-605-150), goat Alexa Fluor 647-coupled anti-rat (1:500, Thermo Fisher Cat#112-005-003) and Streptavidin A488 (1:500, Invitrogen, #S11223) or Streptavidin Atto 550 (1:500, Sigma, #96404). Sections were then rinsed twice in PBS before mounting on SuperFrost® slides (VWR #631-0108) and coverslipped with a xylene-based mounting medium (Leica Micromount #3801731). Slices were stored at 4°C until imaging. Confocal miscroscopy/imaging were carried out at the Montpellier RIO Imaging Facility. Fluorescent images of labeled cells in the region of interest were captured using sequential laser scanning confocal microscopy (Leica SP8). Three to eight images per hemisphere from four to six sections for a given markers were used for quantifications. Adjacent serial sections were never counted for the same marker to avoid any potential double counting of hemisected neurons. Images have been analyzed using Fiji software.

### Tissue collection for polyribosome immunoprecipitation

Whole striata were extracted from adult *Pvalb-Ribotag* mice, as previously described (Puighermanal et al., 2020). Mice chronically treated with d-amph (5 mg/kg) or saline were sacrificed three days after the last administration. Mice that underwent the operant behavior task were euthanized one day after the last FR5 training session. Striatal tissue from two to three mice was pooled to generate a single sample for polyribosome immunoprecipitation and subsequent RNAseq.

### Polyribosome immunoprecipitation

HA-tagged ribosome immunoprecipitation from striatal samples of *Pvalb-Ribotag* mice was performed as previously described (Cutando et al., 2022), using 5 µl of anti-HA antibody (Clone 16B12; BioLegend, 901513) and magnetic beads (Fisher Scientific, #88803). Total RNA from the pellet fractions was extracted using the RNeasy Micro Kit (Qiagen, #73934), and from the input fractions using the RNeasy Mini Kit (Qiagen, #74104), following the manufacturer’s instructions. An on-column DNase treatment was included to eliminate genomic DNA contamination. RNA quality and quantity were assessed using 1 µl of sample on a Nanodrop One spectrophotometer (Thermo Scientific). For the d-amphetamine experiment, three biological replicates were used for RNAseq analysis, each consisting of pooled tissue from 2 to 3 mice. For the operant behavior experiment, six biological replicates were used, each consisting of pooled striata from 2 to 3 mice.

### Low-input RNA sequencing and data processing

RNA sequencing libraries from *Pvalb-Ribotag* mice striatal tissue were prepared following the SMART-seq2 protocol (Picelli et al., 2014), with some modifications. Briefly, total RNA samples were quantified using the Qubit® RNA BR Assay Kit (Thermo Fisher Scientific), and RNA integrity was assessed with the Agilent DNF-471 RNA (15 nt) Kit on the Fragment Analyzer 5200 system (Agilent). Reverse transcription was performed on 1.8 µl of total RNA input (6– 11 ng, depending on sample availability) using SuperScript II (Invitrogen) in the presence of oligo-dT30VN primers (1 µM; 5′-AAGCAGTGGTATCAACGCAGAGTACT30VN-3′), template-switching oligonucleotides (1 µM), and betaine (1 M). The resulting cDNA was amplified using KAPA HiFi HotStart ReadyMix (2×) (Roche) and 100 nM IS PCR primer (5′-AAGCAGTGGTATCAACGCAGAGT-3′), with 8 cycles of PCR. After purification with Agencourt AMPure XP beads (1:1 ratio; Beckman Coulter), product size distribution and concentration were evaluated using the Bioanalyzer High Sensitivity DNA Kit (Agilent). A total of 200 ng of amplified cDNA was fragmented for 10 minutes at 55 °C using the Nextera XT Kit (Illumina), followed by 12 cycles of amplification with indexed Nextera PCR primers. The resulting libraries were purified twice using Agencourt AMPure XP beads (0.8:1 ratio) and quantified again using the Bioanalyzer High Sensitivity DNA Kit. Sequencing was performed on an Illumina NovaSeq 6000 system in paired-end mode with a read length of 2 × 51 bp, according to the manufacturer’s protocol for dual indexing. Image analysis, base calling, and quality scoring were carried out using Real Time Analysis (RTA) software version 3.4.4, followed by the generation of FASTQ files. For the operant behavior experiment (**Fig. 7**), RNA-seq reads were mapped against the *Mus musculus* genome (GRCm38) using STAR 2.5.3a (Dobin et al., 2013) with ENCODE parameters. Gene-level quantification was performed with RSEM 1.3.0 (Li and Dewey, 2011) using the gencode.M21 annotation. For the d-amphetamine experiment (**Fig. 6**), RNAseq reads were aligned against the *Mus musculus* genome (GRCm39) using STAR 2.7.8a (Dobin et al., 2013), also with ENCODE parameters and gene quantification was performed with RSEM 1.3.0 (Dobin et al., 2013) using the gencode.M34 annotation.

### RNAseq differential expression

Genes with at least 1 count-per-million reads (cpm) in at least 3 samples were retained. Counts were normalized using the trimmed mean of M-values (TMM) method and transformed into log2-counts per million (logCPM). Differential gene expression analysis was performed using the limma R package v3.54.2 (Ritchie et al., 2015). The voom function (Law et al., 2014) was used to estimate the mean-variance relationship and compute observation-level weights. These voom-transformed counts were used to fit linear models. Functional enrichment analysis was conducted using g:Profiler via the gprofiler2 R package v0.1.8 (Kolberg et al., 2023) and ShinyGO (Ge et al., 2020). Multidimensional scaling plots were generated with the limma function plotMDS using the top 500 most variable genes.

### Behaviors

#### Locomotor response to d-amphetamine

Mice were tested in a circular corridor system (Imetronic, Pessac, France) for 120 minutes. Horizontal activity was recorded when mice triggered two adjacent infrared beams positioned 1 cm above the floor in each 90° section of the corridor, indicating movement through one-quarter of the circular track. Vertical activity, measured as rearing, was recorded when mice broke beams placed 7.5 cm above the floor, reflecting upright exploratory movements. Prior to drug administration, *Pvalb-Ribotag* mice underwent a two-day habituation period. Each day, they were placed in the activity chamber for 30 minutes, injected with saline, and then returned to the chamber for an additional 90 min. The following days, the same protocol was followed, with the exception that the mice were allocated to two groups: one administered with saline and the other with d-amphetamine (5 mg/kg, i.p.). This paradigm was repeated for five consecutive days.

#### Operant behavior

One week prior the experiments, mice were individually housed. Five days before conditioning, they were food-restricted to maintain 85% of their original body weight. Food restriction was maintained from day 1 to day 9, after which mice had *ad libitum* access to food from day 10 to day 15. Mice were assigned to one of two groups based on the type of food reward used in the operant paradigm. The high-palatable food group received isocaloric pellets (TestDiet) with the same caloric content as standard chow (3.48 kcal/g) but with a higher sucrose content (49% of carbohydrates) and chocolate flavoring. The standard food group received pellets with the same caloric and palatability profile as standard chow. The operant training began with a fixed ratio (FR)-1 schedule of reinforcement, during which mice were presented with two levers. Pressing the active lever resulted in the delivery of a pellet, while pressing the inactive lever had no consequences. Each reward delivery was followed by a 15-s time-out period, which was maintained throughout all experimental phases. Following the FR1 phase, mice underwent four days of FR5 training, where five presses on the active lever were required to receive a pellet. The final phase consisted of 6 additional days of FR5 training, during which mice had *ad libitum* access to food in their home cages. Within each food group, mice were further divided into master and yoked subgroups. Each yoked mouse was paired with a master mouse. Master mice performed the operant task as described, whereas yoked mice received a pellet passively whenever their paired master obtained a pellet. Lever presses by the yoked mice had no consequences.

### Statistical analyses

Statistical analyses were performed with GraphPad Prism v10.4.0. Behaviors were analyzed with two-way repeated measure ANOVA or three-way ANOVA as detailed in Supplemental Table 4.

## Results

### Generation and characterization of *Pvalb-Ribotag* mice

We first generated *Pvalb-Ribotag* mice expressing the channel rhodopsin 2 (ChR2) by crossing *Pvalb^IRES-Cre/+^* mouse line with the *Ai32^loxP^*;*Ribotag^loxP^*line (named thereafter *Pvalb-ChR2-Ribotag*) allowing us to visualize PV interneurons and characterize their translatome. Triple immunofluorescence analyses revealed that endogenous PV interneurons identified using PV antibody outnumbered by far the ChR2- and/or HA-positive interneurons in the DS (**Fig 1a-b**). Indeed, no recombination was found in ∼68% and ∼57% of the PV interneurons analyzed in the dorsomedial (DMS) and dorsolateral striatum (DLS), respectively (**Fig. 1b**). We also found a fraction of PV cells expressing either only ChR2 (DMS: ∼10% and DLS: ∼8%) or to a lesser extend expressing only HA (DMS: ∼1% and DLS: ∼1%) (**Fig. 1-b**). Facing this low rate of recombination, we decided to reevaluate the degree of recombination in two additional mouse lines, the *Pvalb-Ribotag* and the *Pvalb-Ai14* mice expressing the red fluorescent protein tdTomato (**Fig. 1c**). In *Pvalb-Ribotag* mice, double immunofluorescence analyses revealed a higher percentage of PV interneurons expressing HA (**Fig. 1b and d**). We also found that the percentage of PV/HA-expressing interneurons was higher in the DLS (∼75%) compared to the DMS (∼49%). Similar results were found when analyses were performed in *Pvalb-Ai14* mice in which ∼67% and ∼47% of the PV interneurons expressed RFP in the DLS and DMS, respectively (**Fig. 1e-f**). Importantly, in none of the three mouse lines tested, did we detect cells expressing ChR2, HA or tdTomato in the ventral striatum.

**Figure 1:**
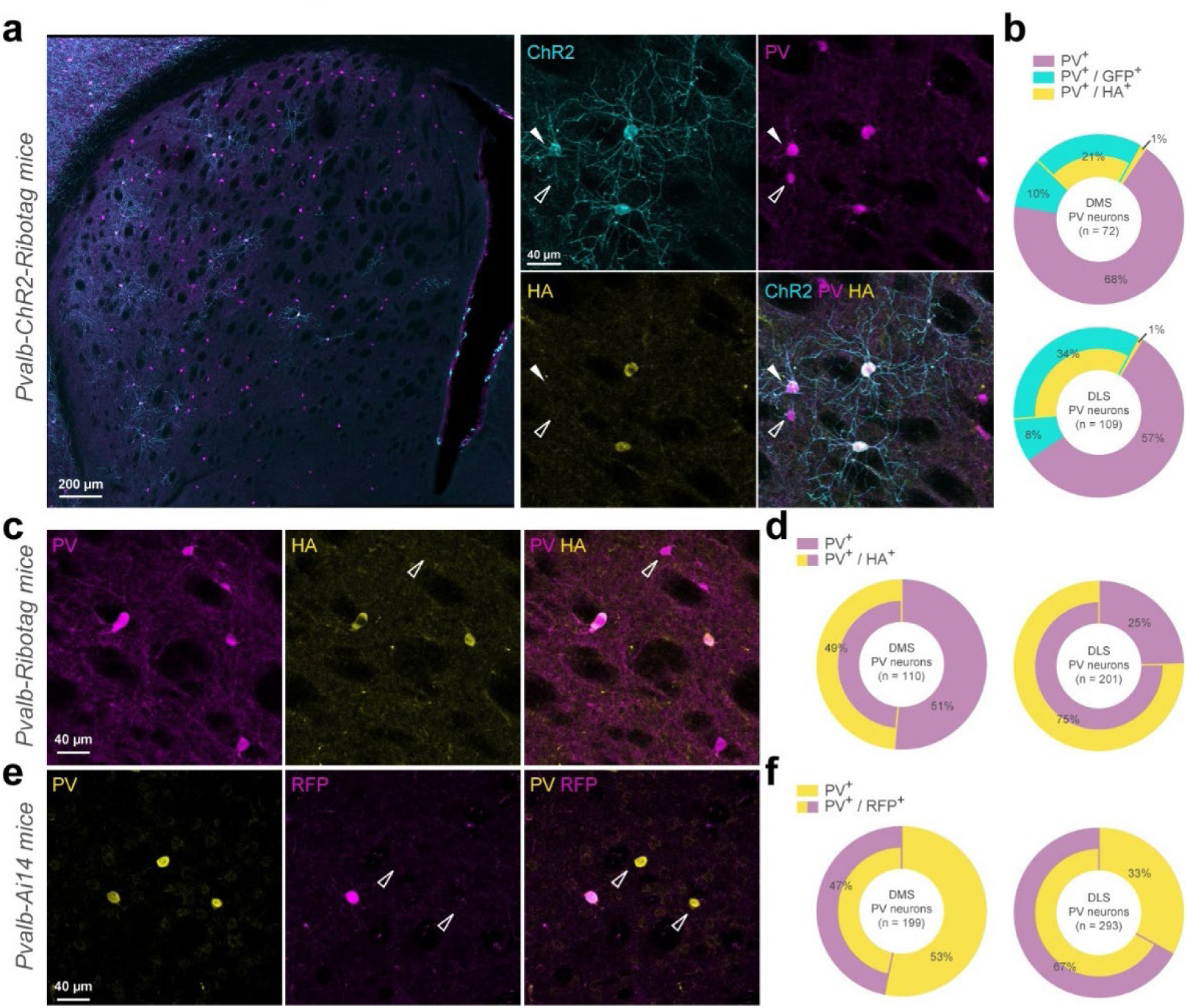
Cre-dependent report proteins expression is restrained to a subpopulation of Parvalbumin-positive interneurons. (**a**) Coronal striatal section from a *Pvalb-ChR2-Ribotag* mouse showing GFP reporter fluorescence (cyan) and stained with PV (magenta), HA (yellow). (**b**) Doughnut charts summarize the proportion of cells expressing either parvalbumin alone (PV+) or combinations of PV+, GFP+ and/or HA+ markers. (**c**) Immunostaining of a *Pvalb-Ribotag* striatal section for PV (magenta) and HA (yellow) and (d) summary data for simple (PV+) or double stained (PV+/HA+) neurons. (**e**) Immunostaining of a *Pvalb-Ai14* striatal section for PV (yellow) and RFP (magenta) and (**f**) summary data for simple (PV+) or double stained (PV+/RFP+) neurons.

### Translatome profile of DS PV interneurons using *Pvalb-Ribotag* mice

The translatome of DS PV interneurons was established by identifying the relative enrichment of genes in the pellet fraction containing tagged Ribosomes-bound mRNAs compared to the input fraction where mRNAs from all cell types were present (**Fig. 2a**). We first validated the selectivity of the approach by demonstrating the enrichment in the pellet fraction of well-established PV markers including *Pvalb, Cox6a2, Kcnc1, Vwc2, Clstn2, Nrip3, Nxph1, Pthlh, Ubash3b* and *Kcnip1* (**Fig. 2b** and **Supplemental Table 1**) (Muñoz-Manchado et al., 2018; Saunders et al., 2018). Conversely, transcripts identifying spiny neurons projections (SPNs: *Gpr88, Ppp1r1b, Arpp19, Pde10a*) from both the direct (dSPNs: *Drd1, Pdyn, Tac1*, *Eya1*) and the indirect pathway (iSPNs: *Gpr6, Adora2a, Penk, Necab1*) (Gokce et al., 2016; Märtin et al., 2019) as well as other classes of striatal GABAergic interneurons (INs: *Npy, Sst, Htr3a, Calb2, Chodl*) and large cholinergic interneurons (CINs: *Chat, Slc17a8, Slc10a4, Ntrk1*) (Muñoz-Manchado et al., 2018) were all decreased (**Fig. 2c** and **Supplemental Table 1**). Similar de-enrichment was found for genes used to classify astrocytes (*Gfap*, *Aldh1l1, S100b*), microglia (*Aif1*, *Trem2*, *Tmem119*) and oligodendrocytes (*Olig2*, *Cnp*, *Mog*) (**Fig. 2d**). Among the 14185 protein-coding genes detected in our RNAseq, 3484 were identified as enriched in DS PV interneurons as compared to the DS inputs (adjusted *p* value of < 0.05) (**Fig. 2e** and **Supplemental Table 1**). This number dropped down to 2750 for those displaying a fold-change > 1.5 (**Fig. 2f** and **Supplemental Table 1**). We also found a significant enrichment of several non-coding transcripts including 663 long non-coding RNAs (LncRNA), 963 To be Experimentally Confirmed RNAs (TEC) and 172 pseudogenes mainly composed of processed pseudogenes (147 out of 172) (**Fig. 2g** and **Supplemental Table 1**).

**Figure 2:**
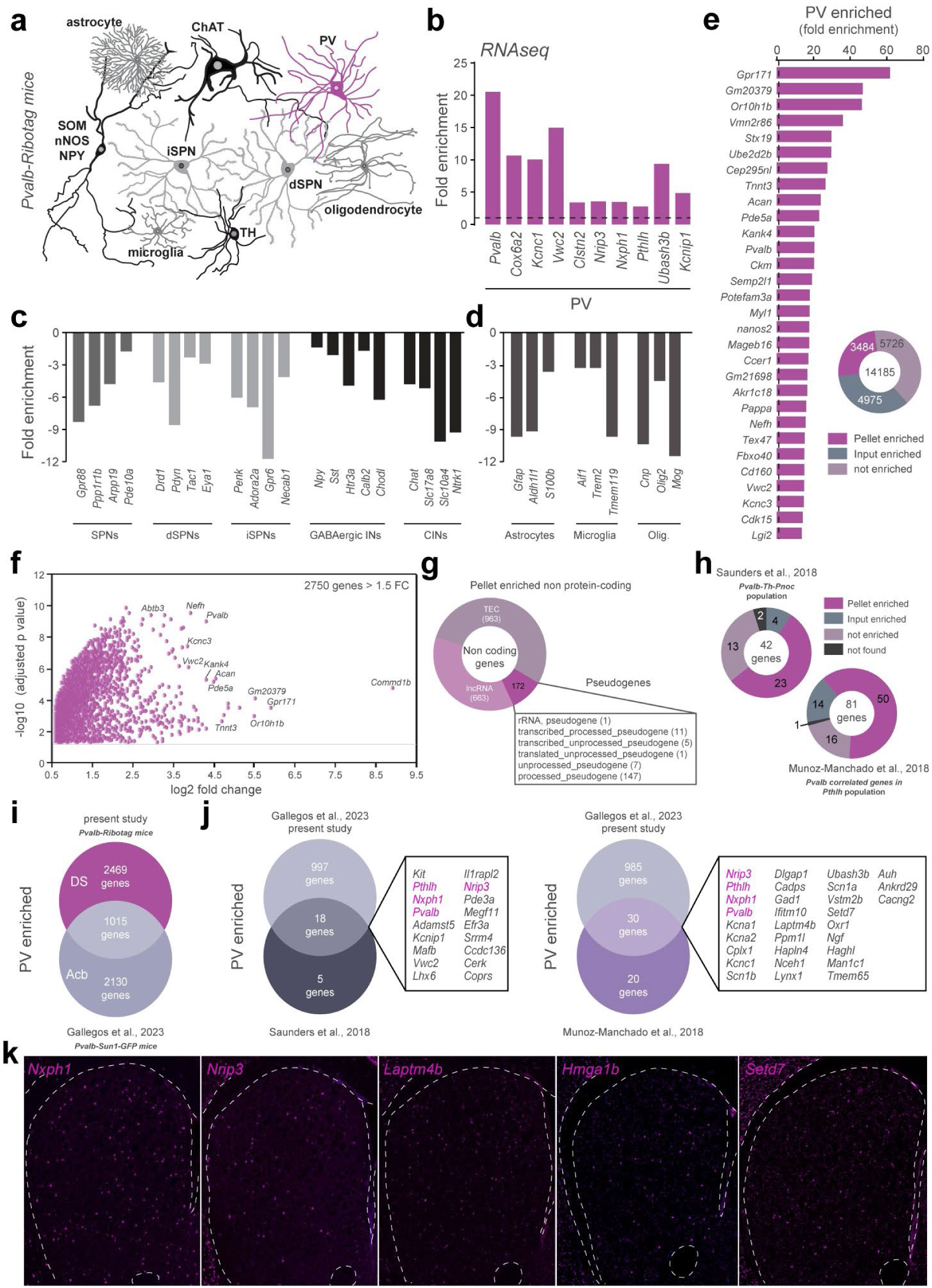
Translatome of PV-positive interneurons using the *Pvalb-Ribotag* mice. (**a**) Cartoon depicting the heterogeneity of striatal cell types. In magenta, the PV-positive cells targeted in *Pvalb-Ribotag* transgenic mice. (**b-d**) Validation by RNAseq of the enrichment of PV interneurons marker genes (**b**) and de-enrichment of markers for other striatal neurons (**c**) and other cell types (**d**) in the pellet fraction from HA-tag pull-down. (**e**) Protein-coding genes enrichment in the pellet fraction. (**f**) Volcano plot of protein-coding genes with a fold-change above 1.5 in the pellet fraction. (**g**) Doughnut plot of non-coding genes relative distribution according to subtypes: pseudogenes, TEC (To be Experimentally Confirmed) and long non-coding RNAs (lncRNA). (**h**) Proportion and number of PV-enriched transcripts previously identified using single cell RNAseq. (**i**) Venn diagram showing the number and overlap of genes enriched in PV interneurons in the dorsal striatum (DS) and nucleus accumbens (Acb). (**j**) Cross-comparison of genes identified in PV interneurons in mouse models from our and 3 different laboratories. The only 4 genes in common (*Pvalb*, *Pthlh*, *Nxph1*, *Nrip3*) are highlighted in purple. (**k**) *In situ* hybridization example images for 5 representative genes expressed in PV interneurons (from the Allen Brain Atlas).

Cross-analyses with previous single-cell RNAseq studies revealed that PV-enriched transcripts found in *Pvalb-Th-Pnoc* (Saunders et al., 2018) and *Pvalb-Pthlh* (Muñoz-Manchado et al., 2018) populations match at 55% and 62% with those identified in the present study (**Fig. 2h** and **Supplemental Table 1**). We also compared the overlap between the genes we found to be enriched in the DS PV interneurons with those enriched in the PV interneurons of the nucleus accumbens (Acb) unveiled using the *Pvalb-Sun1-GFP* mouse line (Gallegos et al., 2023). We identified 1015 shared transcripts enriched between DS and Acb PV interneurons (**Fig. 2i** and **Supplemental Table 1**). Among these common genes, 18 were found in the *Pvalb-Th-Pnoc* population and 30 in the *Pvalb-Pthlh* population (**Fig. 2j** and **Supplemental Table 1**). All in all, only 4 transcripts (*Pvalb*, *Pthlh*, *Nxph1*, *Nrip3*) were found systematically enriched in PV interneurons regardless their location (DS vs Acb) and the RNAseq approaches used (scRNAseq vs TRAP) (**Fig. 2j** and **Supplemental Table 1**). Finally, *in situ* hybridization from the Allen Brain Atlas dataset for *Nxph1*, *Nrip3, Laptm4b, Hmga1b* and *Setd7* transcripts presented a sparse distribution typical of DS PV interneurons (**Fig. 2k**).

### Extracellular matrix and cell adhesion classification of DS PV interneurons

Gene Ontology (GO) enrichment analysis using the ShinyGO v0.8 web tool (Ge et al., 2020) revealed that synapse organization represented one of the most significant GO terms associated to Biological Process and Cellular Component (**Supplemental Fig 1**). Because synaptic stabilization is tightly associated with extracellular matrix (ECM) structures (Christensen et al., 2021; Wingert and Sorg, 2021), we examined whether ECM-related genes were enriched in DS PV interneurons using the MatrisomeDB database. Among 274 annotated core matrisome genes, we found transcripts encoding ECM glycoproteins (19), collagens (6) and proteoglycans (5) enriched in PV interneurons (**Fig. 3a-b** and **Supplemental Table 2**). We also identified several matrisome-associated genes among which some encode ECM regulators (22), ECM-affiliated proteins (12) and secreted factors (31) (**Fig. 3a-b** and **Supplemental Table 2**). Interestingly we noticed that 44 ECM-related genes enriched in DS PV interneurons were also found in Acb PV interneurons (labelled in magenta) (**Fig. 3a** and **Supplemental Table 2**). Closed inspection of the identity of proteoglycan-enriched genes revealed the presence of core constituents of perineuronal nets (PNNs) including *Acan* and *Vcan* transcripts encoding for two chondroitin sulfate proteoglycans (Aggracan and Versican) and *Hapln1* and *Hapln4* encoding for hyaluronan and proteoglycan link protein 1 and 4 (**Fig. 3a** and **Supplemental Table 2**). To examine the presence of PNNs around DS PV interneurons, we performed Wisteria floribunda agglutinin (WFA) staining (Härtig et al., 2022) on striatal slices from *Pvalb-ChR2*, *Pvalb-Ai14* and *Pvalb-Ribotag* mice in which PV interneurons were identified indirectly with GFP, tdTomato or HA and directly using PV antibody (**Fig. 3c**). Immunofluorescence analysis revealed that regradless the mouse line and PV detection method used, PNNs ensheath the majority of the DS PV interneurons, with WFA staining observed in more than 94% of them (**Fig. 3c**).

**Figure 3:**
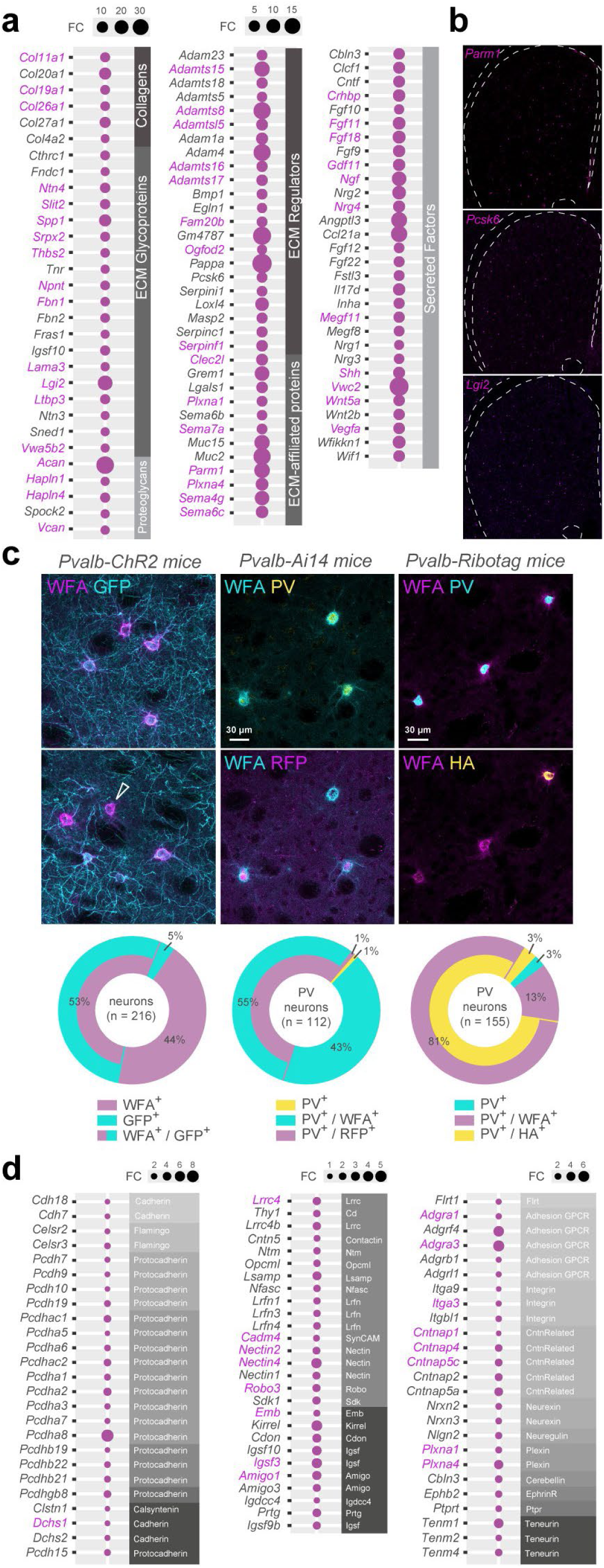
Molecular signature of DS PV interneurons transcripts related to extracellular components. (**a**) Extracellular matrix (ECM) genes enriched in DS PV interneurons, classified by sub-categories (grey bands). FC, fold change. Magenta, genes also enriched in Acb PV neurons (**b**) *In situ* hybridization example images for 3 representative ECM genes expressed in PV interneurons (Allen Brain Atlas). (**c**) Perineuronal nets identified using Wisteria floribunda agglutinin (WFA) in the 3 different *Pvalb* reporter mouse lines used in this study, highlighting the presence of ECM around the majority of PV-positive cells and only a minority of PV-negative cells (arrow). Doughnut charts depicting the percentage of co-labelling for the different reporters. (**d**) Cell adhesion molecules transcripts enriched in DS PV interneurons. Enriched genes in common between DS and Acb are highlighted in magenta.

Cell adhesion molecules are also core constituents of synaptic organization and stabilization. We therefore analyzed the distribution of PV-enriched genes among cadherins, protocadherins, integrins as well as several different categories of cell adhesion molecules (**Fig. 3d** and **Supplemental Table 2**). We found that genes encoding alpha isoforms of clustered protocadherins (9 out of 13: *Pcdhac1, Pcdhac2, Pcdha5, Pcdha6, Pcdha1, Pcdha2, Pcdha3, Pcdha7, Pcdha8*), non-clustered protocadherins (5 out of 12: *Pcdh7*, *Pcdh9, Pcdh10, Pcdh19, Pcdh15*), contactin-associated proteins (5 out 7: *Cntnap1*, *Cntnap4*, *Cntnap5c*, *Cntnap2*, *Cntnap5a*) as well as amigo (2 out of 3: *Amigo1*, *Amigo3*) or neurexin (2 out of 3: *Nrx2*, *Nrx3*) were particularly enriched in DS PV interneurons (**Fig. 3d** and **Supplemental Table 2**). In contrast to ECM-related genes, only 17 cell adhesion-related genes were common to DS and Acb PV interneurons (labelled in magenta). These findings allow the identification of a unique ECM and adhesion molecule signature for DS PV interneurons (**Fig. 3d** and **Supplemental Table 2**).

### Neurotransmitter system and ion channels classification of DS PV interneurons

Cellular Component GO terms associated with synapses functions were also highly represented in our analysis (**Supplemental Fig 1**). We therefore took advantage of a system classification previously implemented allowing the rapid visualization of genes encoding receptors, transporters, and enzymes involved in the turnover of a given neurotransmitter system including serotonergic, catecholaminergic, GABAergic, cholinergic and glutamatergic (Puighermanal et al., 2020) (**Fig. 4a-e** and **Supplemental Table 3**). These analyses revealed the enrichment of a restricted set of serotonergic (*Htr2b*, *Htr7*) (**Fig. 4a** and **Supplemental Table 3**) and catecholaminergic (*Drd4*, *Adra1a*) receptors (**Fig. 4b**). Several components of the GABAergic system were also found to be enriched including genes encoding GABAa and GABAb receptor subunits (*Gabra1*, *Gabrg3, Gabrd, Gabbr2*) as well as transcripts encoding proteins involved in GABA turnover (*Gad1*, *Gad2*) and transport (*Slc6a1*, *Slc6a8*) (**Fig. 4c** and **Supplemental Table 3**). *Chrnb2* was the only cholinergic gene found to be enriched in DS PV interneurons (**Fig. 4d**). Regarding the glutamatergic system, 10 out of the 18 genes encoding ionotropic glutamate receptors were enriched in PV interneurons (*Grik1, Grik3, Grik5, Gria3, Gria4, Grin1, Grin2a, Grin2b, Grin2d, Grid2*) while none of the transcripts encoding metabotropic glutamate receptors (*Grm*) were found to be enriched (**Fig. 4e** and **Supplemental Table 3**). Finally, we found that DS and Acb PV interneurons only shared 5 neurotransmitter system-related genes, 3 associated to the GABAergic system (*Gabra1, Gad1, Slc6a8*) and 2 with the glutamatergic one (*Gria4, Grin2d*) (labelled in magenta) (**Fig. 4**).

**Figure 4:**
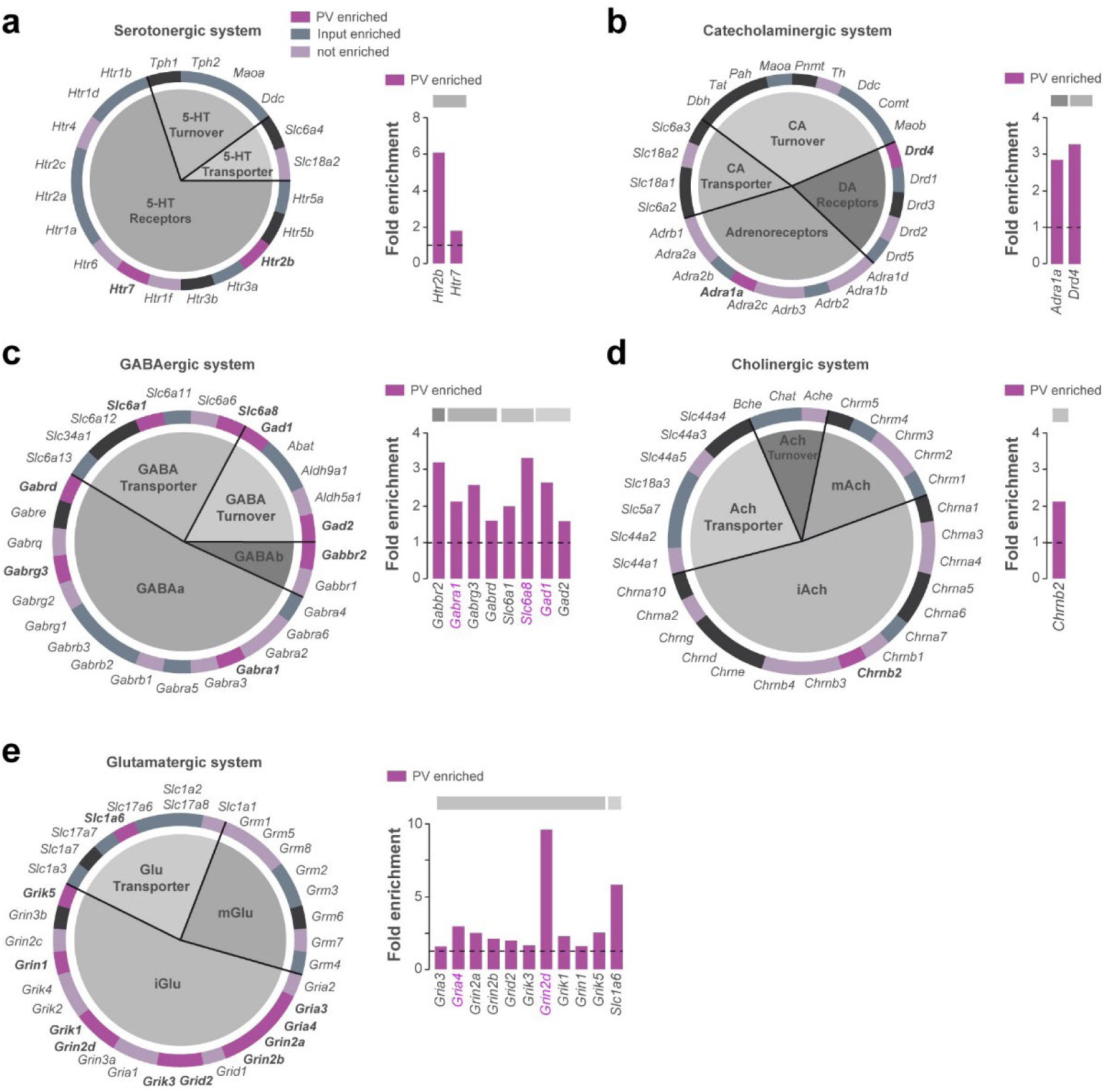
Neurotransmitter systems-related genes enriched in DS PV interneurons. Pie chart and fold enrichment graphs of DS PV genes within the (**a**) serotoninergic, (**b**) catecholaminergic, (**c**) GABAergic, (**d**) cholinergic and (**e**) glutamatergic systems. In dark purple are highlighted the genes enriched in the pellet, in grey the ones enriched in the inputs, in light purple the ones detected but not enriched and in black those not found in either the pellet or the input. Enriched genes in common between DS and Acb are highlighted in magenta.

We also performed a complete classification of voltage-gated ion channels (**Fig. 5** and **Supplemental Table 3**), including calcium channels (**Fig. 5a**), sodium channels (**Fig. 5b**), and potassium channels comprising voltage-gated potassium channels (**Fig. 5**), inwardly rectifying potassium channels (**Fig. 5d**), two-P potassium channels (**Fig. 5e**), calcium-activated potassium channels (**Fig. 5f**), and accessory subunits (**Fig. 5g**). The main pore-forming alpha 1 subunits of voltage-gated calcium channels were particularly enriched compared to the other subunits (7 out of 10 constituents) (**Fig. 5a**). Voltage-gated potassium channels distributed among the different families were also highly enriched in DS PV interneurons, several of them displaying a fold-enrichment > 5 (*Kcna2, Kcnc1, Kcnc3, Kcng4, Kcnh2*) (**Fig. 5c**). Our analysis also unveiled a biased expression in DS PV interneurons of G-protein activated inward rectifying potassium channels (*Kcnj3, Kcnj6, Kcnj9, Kcnj11, Kcnj12*) (**Fig. 5d**). In contrast to neurotransmitter system-related genes, the expression of voltage-gated ion channels appears to be more conserved between DS and Acb PV interneurons as suggested by the important shared enriched expression of genes encoding voltage-gated potassium channels (9 out of 13 genes including *Kcna1, Kcna2, Kcnc1, Kcnc2, Kcnc3, Kcnb2, Kcng4, Kcns3, Kcnh2*) (**Fig. 5c**), G-protein gated potassium channels (3 out of 5 genes: *Kcnj3, Kcnj9, Kcnj12*) (**Fig. 5d**), Two-P potassium channels (*Kcnk3, Kcnk12*) and accessory subunits (3 out 3 genes: *Kcnab2, Kcnab3, Kcnmb2*) (**Fig. 5g**).

**Figure 5:**
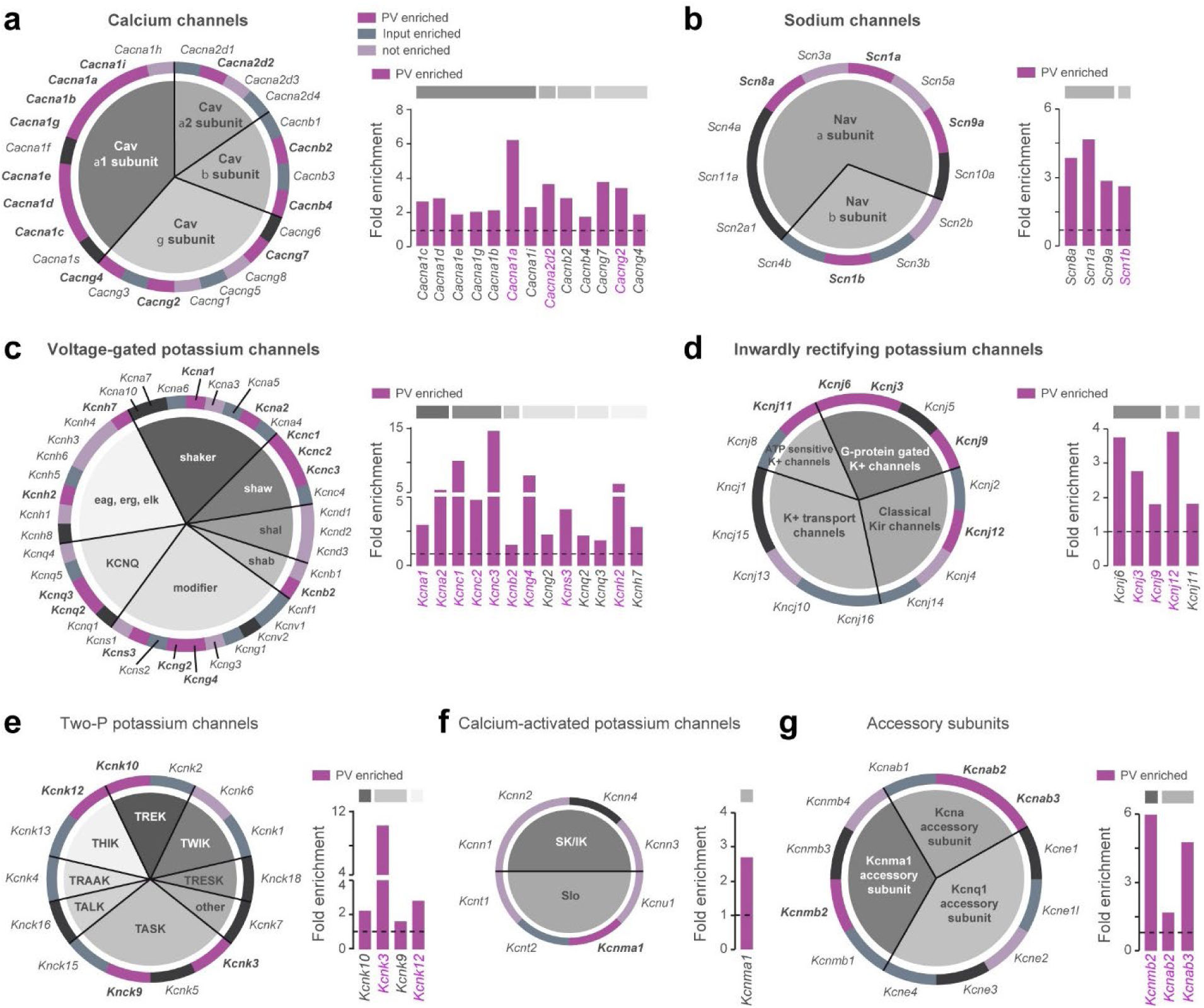
Ion channel genes enriched in DS PV interneurons. Pie chart and fold enrichment graphs of DS PV genes encoding for (**a**) calcium channels, (**b**) sodium channels, (**c**) voltage-gated potassium channels, (**d**) inwardly rectifying potassium channels, (**e**) two-P potassium channels, (**f**) calcium-activated potassium channels and (**g**) channels’ accessory subunits. In dark purple are highlighted the genes enriched in the pellet, in grey the ones enriched in the inputs, in light purple the ones detected but not enriched and in black those not found in either the pellet or the input. Enriched genes in common between DS and Acb are highlighted in magenta.

**Figure 6:**
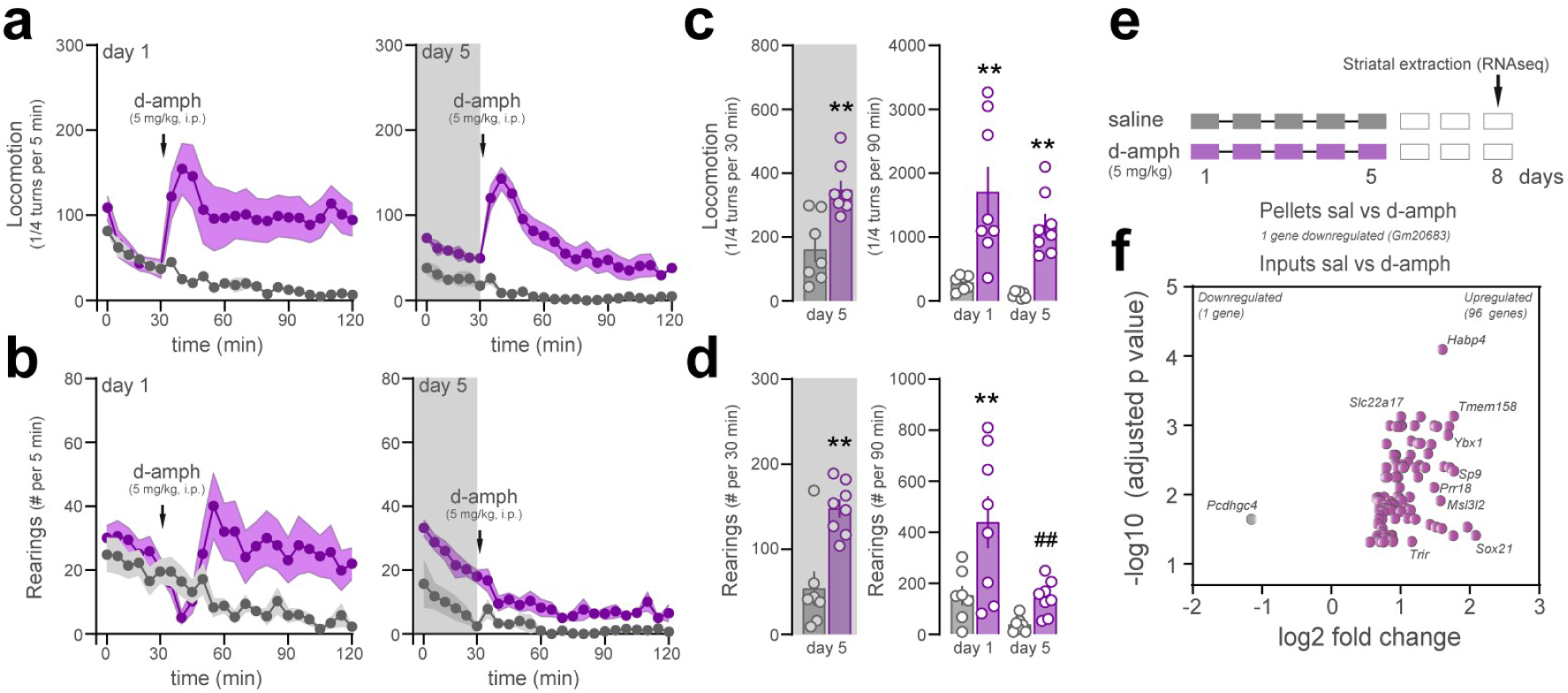
Limited gene expression changes induced in PV interneurons by repeated psychostimulant administration. (**a**) Locomotor activity (top) and (**b**) rearings (bottom) induced by d-amphetamine on day 1 (left) and day 5 (right) of administration (5 mg/kg i.p., purple; saline controls, grey). (**c-d**) Summary histograms showing locomotion (**c**) and rearings (**d**) on day 5 before drug administration (left, grey area) and comparison between day 1 and day 5 locomotion and rearings (right). (**e**) Administration protocol showing RNAseq analysis being performed 8 days after the first d-amphetamine (or saline) injection. (**f**) Volcano plot of significantly enriched or de-enriched genes in total striatal tissue (input) upon d-amphetamine treatment. One downregulated and 96 upregulated transcripts were identified. Of notice, only one gene (*Gm20683*) was found downregulated by d-amphetamine specifically in PV interneurons (pellet). ** p < 0.01 saline vs. d-amph; ## < 0.01 day 1 vs. day 5 (detailed statistical analysis in Supplemental Table 4: 6a-d).

**Figure 7:**
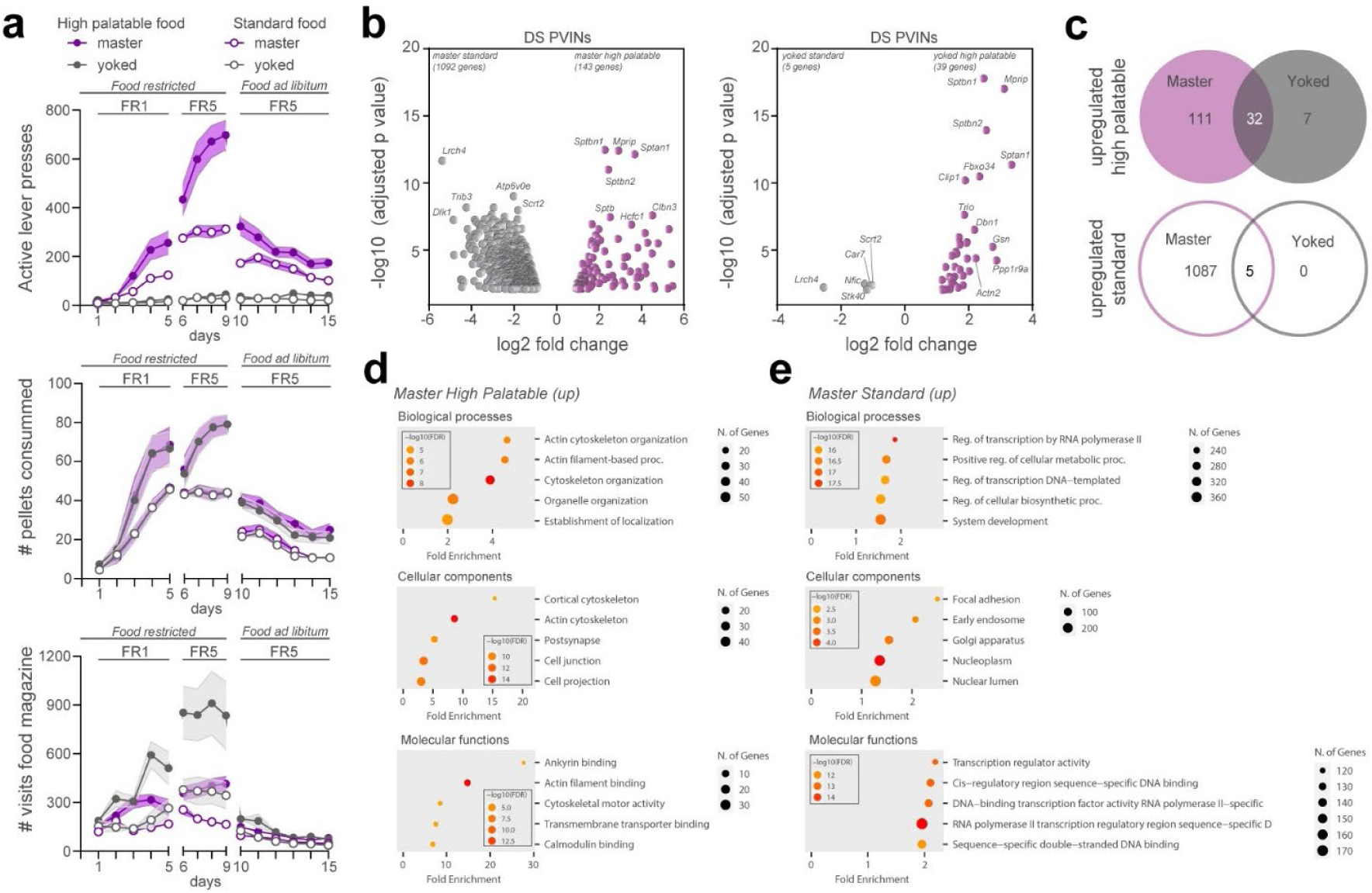
Gene expression changes induced in PV interneurons by food-seeking behaviors. (**a**) Operant behavioral task performance in master (purple) and yoked (grey). Number of active lever presses, pellet consumed and visits to the distributor compartment (magazine) are shown for highly palatable (filled symbols) or standard food (empty symbols) during simple (fixed ratio 1, FR1) and more demanding tasks (FR5) in food restricted animals and upon reintroduction of ad libitum food. (**b**) Volcano plots of significantly enriched or de-enriched genes in DS PV interneurons of master (left) and yoked (right) mice one day after the last session of food-seeking behavior. (**c**) Venn diagrams showing the number of overlapping upregulated genes in master and yoked mice under highly palatable and standard food diet. (**d-e**) Gene ontology analysis of the biological processes, cellular components and molecular functions of the upregulated genes in master animals under the two food conditions. (detailed statistical analysis in Supplemental Table 4: 7a-c)

### Transcription factors classification of DS PV interneurons

Finally, to gain insights into the specification of DS PV interneurons, we classified transcription factors by family using the AnimalTFDB v4.0 database (Shen et al., 2023) (**Supplemental Fig 2**). We analyzed the distribution of DS PV interneurons enriched genes among 6 core families of transcription (**Supplemental Fig 2**). The classification was further refined by analyzing the distribution DS PV neurons enriched genes within each transcription factors subfamilies. The filtering of enriched genes for a fold-change > 4 allowed the description of a putative transcription factors landscape of DS PV interneurons comprising 5 members of the basic domain groups (*Tfap2b, Hes3, Bach2, Maf, Mafb*) (**Supplemental Fig 2a**), 1 member of beta scaffold factors (*Lin28b*) (**Supplemental Fig 2b**), 8 transcripts from the helix-turn helix group (*Onecut2, Satb1, Arid3a, Arx, Barx2, Lhx1, Six4, Pou5f2*) (**Supplemental Fig 2c**), 4 genes from other alpha helix groups (*Sox12, Sox6, Tox2, Tox3*) (**Supplemental Fig 2d**), 5 transcripts from the unclassified structure (*Hmga1b, Pura, Prug, Zfp318, Zfp750*) (**Supplemental Fig 2e**) and 11 members of the zinc coordinating group (*Esrrb, Essrg, Nr6a1, Zfp618, Zim1, Gm14308, Gm14408, Gm14443, Prdm11, Scrt1, Scrt2*) (**Supplemental Fig 2f**). Among all the genes encoding transcription factors enriched in the DS PV neurons, 75 transcripts were also found in the Acb PV neurons dataset (colored transcripts) with percentages of overlap ranging from 40% for the basic domain groups (8 out of 20 genes in common) and 35% for the helix-turn helix group (20 out of 57 common transcripts) to 12.5% for the beta scaffold factors (1 gene out of 8 genes) (**Supplemental Fig 2a-c**).

### Translatome of DS PV neurons upon repeated d-amphetamine exposure

Previous works showed that striatal PV interneurons were required for the action of psychostimulant drugs (Wang et al., 2018; Wiltschko et al., 2010). We therefore sought to determine the impact of repeated exposure to d-amphetamine on the DS PV interneurons translatome. *Pvalb-Ribotag* mice used for RNAseq were administered either saline or d-amphetamine (5 mg/kg, i.p.) for 5 consecutive days during which psychomotor responsiveness was measured in a circular corridor (**Fig. 6**). As expected, d-amphetamine-treated mice exhibited a robust increase in both horizontal (locomotion) and vertical (rearings) locomotor activity as compared to saline-treated mice (**Fig. 6a**). Although repeated d-amphetamine administration triggered a robust conditioned locomotor responses to the context, mice failed to develop locomotor sensitization and strongly decreased their rearing behaviors at the expense of the development of stereotyped behaviors (**Fig. 6b-d**). To detect stable modifications, tagged ribosomes-bound mRNAs from DS PV interneurons were isolated 3 days after the last administration of d-amphetamine and compared to DS PV interneurons isolated mRNAs from saline-treated *Pvalb-Ribotag* mice (**Fig. 6e**). Comparison between PV neurons saline and d-amphetamine samples identified only one differentially translated transcripts (*Gm20683*) between these groups (**Fig. 6e**). In contrast, when input fractions from saline and d-amphetamine samples were compared, we identified 1 downregulated and 96 upregulated transcripts in d-amphetamine group most of them being microtubule cytoskeleton-related genes (**Fig. 6f**).

### Translatome of DS PV neurons following standard and high palatable food-seeking behaviors

To evaluate whether reward type and contingency impacted gene expression in DS PV neurons, food-restricted *Pvalb-Ribotag* mice were divided into two groups based on pellet type: a high-palatable diet or a standard chow (Castell et al., 2024). Within each dietary condition, animals were subsequently allocated to two groups: master mice, which earned food rewards through lever pressing, and yoked mice, which passively received a pellet each time their paired master obtained one. During the initial FR1 schedule (days 1 to 5), behavioral performance began to diverge after the third training session (**Fig. 7a, upper panel**). Compared to master mice given regular chow, master mice trained for high-palatable rewards exhibited a significantly higher number of active lever presses (**Fig. 7a, upper panel**). Accordingly, both high-palatable masters and their yoked counterparts received a greater number of pellets than the standard food groups (**Fig. 7a, middle panel**). As expected, lever pressing remained low and stable across days in yoked animals, reflecting the absence of action-outcome contingency (**Fig. 7a, upper panel**). Upon transition to an FR5 schedule (days 6–9), high-palatable masters showed a progressive increase in lever pressing across sessions. In contrast, standard food masters exhibited a marked increase in responding on the first day of FR5, followed by performance stabilization, as reflected in both lever press frequency and pellet delivery (**Fig. 7a, upper and middle panels**). From day 10 to 15, animals were returned to *ad libitum* feeding. Under these conditions, both groups of master mice reduced operant responding, consistent with reduced motivational drive as their physiological needs were now met. High-palatable masters continued to lever press at a higher rate than standard chow masters, although both groups demonstrated a progressive decline in responding over time (**Fig. 7a, upper panel**). To further highlight the behavioral dissociation between master and yoked groups, we quantified food magazine visits (**Fig. 7a, lower panel**). During the food-restricted phase, yoked mice exhibited a higher frequency of magazine checks compared to their corresponding master mice consistent with the non-contingent nature of reward delivery in yoked animals. This pattern was abolished under ad libitum feeding, during which all groups displayed comparable low levels of food magazine inspection (**Fig. 7a, lower panel**). One day after the last session, immunoprecipitation of tagged ribosomes was performed and isolated DS PV interneurons mRNAs from the 4 groups were analyzed by high-throughput RNAseq. Paired comparisons between master groups (standard vs high palatable) revealed the highest number of the differentially translated transcripts (**Fig. 7b** and **Supplemental Table 2**). Indeed, 1092 genes were found to be enriched in DS PV interneurons in standard food masters whereas 143 genes were more expressed in high palatable food masters (**Fig. 7b**). In contrast, analysis between yoked groups (standard vs. high palatable food) identified 44 genes displaying differential expression DS PV interneurons (**Fig. 7b**). However, these latter changes were unlikely directly linked to food palatability as 37 genes out of 44 were similarly modified between standard and high palatable master mice, suggesting that most of the changes observed in DS PV interneurons were a snapshot of the interaction between food-seeking behavior and the palatability (**Fig. 7b**). Comparison of upregulated genes in master and yoked mice fed a high-palatable diet revealed 32 genes commonly upregulated in both groups, indicating that these genes may be associated with the sucrose content of the diet rather than operant behavioral contingency (**Fig. 7c**). In contrast, only 5 genes were commonly upregulated between master and yoked mice on the standard diet, suggesting a more limited shared transcriptional response under these conditions (**Fig. 7c**). To get insights into the biological functions of the changes observed we conducted GO enrichment analysis. We found that most of the genes upregulated in high palatable food master mice were related to actin cytoskeleton organization (**Fig. 7d**). Although transcripts related to tubulin, actin, cell adhesion and ECM regulation were also increased in standard master mice, GO terms associated to transcription were the most represented in this group (**Fig. 7d-e**). Among the 1092 upregulated genes, 155 were identified as transcription factors distributed in the 6 core families.

## Discussion

The present work provides a comprehensive analysis of translated mRNAs enriched in the PV interneurons of the dorsal striatum. Our approach using *Pvalb-Ribotag* mice allowed the identification of more than 2,700 protein-coding transcripts with a fold-change > 1.5 enriched in DS PV interneurons. Our study extends previous single-cell RNAseq works revealing the molecular heterogeneity of striatal PV interneurons (Gallegos et al., 2023; Muñoz-Manchado et al., 2018; Saunders et al., 2018). Finally, our work also unveiled the potential limitation of using the *Pvalb-IRES-Cre* mouse line for studies aiming to investigate the role of PV interneurons across both the dorsal and ventral striatum.

Since the early 2000s, the *Pvalb-IRES-Cre* mouse line has been extensively used to study the role and function of striatal PV interneurons. In most cases, this has been achieved by transducing engineered viral vectors selectively in PV interneurons allowing to establish a causal link between their activity and diverse striatal-dependent behaviors including choice execution (Gage et al., 2010), learning strategies (Owen et al., 2018), habits formation (O’Hare et al., 2017) as well as early phase of reward conditioning (Lee et al., 2018). We therefore decided to use the *Pvalb-IRES-Cre* mouse line to generate *Pvalb-Ribotag* mice with the aim to perform the in-depth analysis of the translatome of striatal PV interneurons. However, during the characterization of this mouse line, we were surprised by the unexpectedly low rate of recombination. Indeed, despite the fact that the *Pvalb-IRES-Cre* mouse line is a knockin to the endogenous *Pvalb* promoter/enhancer elements (Hippenmeyer et al., 2005), we were unable to detect any labeling in the ventral striatum. In the DS the expression of HA was restricted to ∼60% of endogenous PV interneurons with a higher level of recombination in the DLS compared to the DMS reminiscent to the medio-lateral gradient of DS PV interneurons (Berke et al., 2004; Fino et al., 2018). Because the ability of the Cre line to recombine the reporter transgene depends on the nature of that transgene, we initially thought that low recombination efficiency was due to the Ribotag allele not being inserted in the ROSA locus (Sanz et al., 2009). However, we rapidly ruled out this hypothesis as we found a similar rate of recombination with two reporter mouse lines carrying the tdTomato (Ai14) or the channelrhodopsin (Ai32) both inserted in the ROSA locus (Madisen et al., 2012, 2010). An alternative hypothesis would be that in the subset of striatal interneurons expressing low levels of PV, the threshold of Cre expression required to recombine the transgenes is not met, thereby resulting in a large proportion of PV interneurons remaining unlabeled (Liu et al., 2013). Our results extend previous observations of a similarly low rate of recombination in the perirhinal cortex (Nigro et al., 2021). Future studies using the newly developed AAV toolbox enabling to target striatal PV interneurons with high precision should overcome this limitation (Hunker et al., 2025).

Despite this limitation, we successfully trapped HA-tagged cells allowing us to establish the translatome of this restricted population of DS PV interneurons. Comparison of our results with previous datasets suggested that our translatome most likely overlap with the *Pvalb-Pthlh* sub population (Muñoz-Manchado et al., 2018). Moreover, our cross-analysis with the translatome of the Acb identified 1015 transcripts enriched in striatal PV interneurons irrespective of their location (Gallegos et al., 2023). This number, representing about 30% of the PV-enriched genes in both DS and Acb suggests that PV interneurons display a high level of molecular heterogeneity throughout the dorso-ventral axis of the striatum, an observation reminiscent to the one previously reported for the translatomes of *Drd1*- and *Drd2*-SPNs (Montalban et al., 2022; Puighermanal et al., 2020). It will be interesting to determine in the future to what extent the differences observed between the Acb and DS PV interneuron translatomes are functionally relevant and potentially related to the distinct developmental origins of PV interneurons (Knowles et al., 2021).

The use of several databases allowed us to refine our analysis based on systematic functional gene classifications. For instance, the identification of a unique ECM and adhesion molecule signature for DS PV interneurons provides a strong molecular rationale for the presence of perineuronal nets enwrapping PV interneurons (Santos-Silva et al., 2024). Moreover, our system-based classification afforded important insights regarding their electrophysiological signatures. Thus, our analysis revealed the enrichment genes encoding various classes of ion channels previously identified to i) facilitate their rapid repolarization and sustained fast-spiking activity (*Kcna1, Kcna2, Kcnc1,* and *Kcnc2*) (Hu et al., 2014), ii) maintain their resting membrane potential and regulate their excitability (*Kcnj3, Kcnj6, Kcnj11*, and *Kcnj12*) (Du et al., 1996; Furdan et al., 2025; Muñoz-Manchado et al., 2018). Interestingly, the transcripts that support these electrophysiological features are enriched in both DS and Acb PV interneurons, representing a genetic portfolio linked to the function rather than the location of these cells. DS PV interneurons also exhibited marked upregulation of several transcripts encoding calcium (*Cacna1a*, *Cacng7*, *Cacng2, Cacna2d2*) and sodium (*Scn1a*, *Scn8a*, *Scn9a, Scn1b*) channels underlying their electrophysiological properties (Ferguson et al., 2023; Lupien-Meilleur et al., 2021; Rendón-Ochoa et al., 2018). However, it is important to keep in mind that our dataset most likely reflects the translatome of PV interneurons located in DLS and that probably a different, but partly overlapping, set of ion channels may explain some of the electrophysiological features of DMS PV interneurons (Koós and Tepper, 1999; Monteiro et al., 2018). We also identified enriched transcripts that have been previously associated with important functions of PV interneurons in other brain areas. For instance, DS PV interneurons are enriched in *Drd4* and *Errb4* transcripts which encode the dopamine D4 receptor and the neuregulin receptor respectively, providing a potential mechanism through which dopamine could modulate gamma oscillations as previously shown in the hippocampus (Andersson et al., 2012). Finally, this dataset classification also represents an invaluable resource to interrogate whether pathological conditions associated to gene-related disorders could be ascribed to a dysfunction of PV interneurons. Thus, motor symptoms associated to *GRIN2D*- and *SLC6A1*-related disorders including dystonia, chorea and hyperactivity and tremor among others (Cha et al., 2025; Chiu et al., 2005; Gawande et al., 2023; Goodspeed et al., 2020, 1993; Vinnakota et al., 2023), could result from dysfunction of these genes encoding the glutamate ionotropic receptor NMDA subunit 2D (*Grin2d*) and the GABA transporter (*Slc6a1*) respectively, which are both highly enriched in DS PV interneurons. In the same line, motor stereotypies associated to the invalidation of the PV enriched gene *Cntnap2* have been causally linked to hyperexcitability of striatal PV interneurons (Thabault et al., 2024). Finally, mice lacking the creatine transporter *Slc6a8* (3-fold enriched in our dataset) selectively in PV interneurons recapitulated numerous features of the Creatine Transporter Deficiency (CTD), an X-linked neurometabolic disorder presenting with intellectual disability, autistic-like features, and epilepsy (Ghirardini et al., 2023). Future studies can use the same strategy to determine whether other PV-enriched genes identified in our dataset are critical for the normal function of DS PV interneurons and if their deregulation is central in disease pathogenesis.

Previous works indicate that striatal PV interneurons are required for d-amphetamine effects. Thus, DS PV interneurons increased their firing rate in response to a single d-amphetamine administration, an electrophysiological response positively correlated with increased locomotion (Wiltschko et al., 2010). The silencing of Acb PV interneurons impaired d-amphetamine-induced psychomotor sensitization and conditioned place preference (Wang et al., 2018). Finally, important transcriptional regulations have been identified in Acb PV interneurons in response to acute and repeated d-amphetamine exposure (Gallegos et al., 2023). Surprisingly, our analysis identified only 1 transcript (*Gm20683*) differentially regulated in DS PV interneurons of mice repeatedly administered with d-amphetamine compared to saline-treated mice. Apart from the regions analyzed (DS vs. Acb), other factors including the experimental design could account for these differences. Indeed, Gallegos and colleagues injected d-amphetamine at a dose of 3 mg/kg for 7 days, while in our case mice were administered with 5 mg/kg of d-amphetamine once daily during 5 days. Moreover, the different delay between the last administration of d-amphetamine and the extraction of samples (24 hours against 3 days in our protocol) might also explain the lack of differentially regulated genes in our RNAseq.

Besides this, we identified long-lasting changes in the translatome of DS PV interneurons in mice trained to obtain standard or high palatable food pellets compared to yoked-groups receiving them passively. These results suggest that alterations in gene expression in DS PV interneurons may occur under specific conditions, notably during behavioral paradigms requiring optimized execution of sequential motor plans, such as the FR5 reinforcement schedule, in which mice pressed the lever more than 700 times on average to obtain highly palatable rewards. Interestingly, among the upregulated genes, several encode for cell adhesion molecules and cytoskeleton proteins necessary for the function of PV interneurons synapses. Thus, remodeling of DS PV interneurons translatome could contribute to the plasticity underlying the rearrangement of striatal network activity required to optimize motor plans.

Overall, our findings provide a comprehensive molecular resource to study dorsal striatal PV interneurons and highlight their dynamic translatome as a potential substrate for experience-dependent plasticity relevant to both motor function and neuropsychiatric disorders.

## Acknowledgements

The authors thank all the lab members and the Microscopy Imaging Platform from IGF and INM (Biocampus). The authors also thank iExplore at IGF for their involvement in the maintenance and breeding of the colonies. This work was supported by CNRS, INSERM, Fondation pour la Recherche Médicale (EQU202203014705, EV), the French National Research Agency (ANR-20-CE14-0020, ANR-21-CE16-0028, ANR-16-CE16-0018 to EV). L. Cutando was supported by MINECO (Ramon y Cajal) fellowship (Spain) (RYC2022-037332-I), H2020-MSCA-IF-2020 (Proposal 101028078; MITORett) and the postdoctoral Labex EpiGenMed fellowship (Investissements d’avenir, ANR-10-LABX-12-01). CNAG acknowledges the support of the Spanish Ministry of Science and Innovation through the Instituto de Salud Carlos III and the 2014–2020 Smart Growth Operating Program, co-financed with the European Regional Development Fund (MINECO/FEDER, BIO2015-71792-P). We also acknowledge the support of the Generalitat de Catalunya through the Departament de Salut, the Departament de Recerca i Universitats and the Departament d’Empresa i Coneixement.

